# Predicting residue ionization of OmpF channel using Constant pH Molecular Dynamics as benchmarking

**DOI:** 10.1101/2025.06.16.659857

**Authors:** Ernesto Tavares-Neto, Marcel Aguilella-Arzo, Vicente M. Aguilella

## Abstract

Electrostatic interactions play a key role in protein structure function. There is a large family of mesoscopic protein channels whose selectivity is mainly controlled by the protein electrostatic properties and ion specific channel interactions play a minor role. The knowledge of the charge state of the ionizable residues over a wide pH range, often summarized in their pKa, stands as the most valuable information for structure-function studies of many protein channels. However, experimental pKa determination is a difficult task, typically accomplished using Nuclear Magnetic Resonance only in a limited number of membrane proteins. Thus, the pKa calculation is the most frequently used alternative. Constant pH Molecular Dynamics (CpHMD) simulation provides arguably the most accurate pKa prediction method in proteins containing many charged residues since it captures the coupling between conformational dynamics and residue protonation. Here we study the charge state of a general diffusion porin, OmpF, in which protons exert a crucial regulation of the channel discrimination of small inorganic ions as well as antibiotic translocation. We examine the pKa prediction using different methods, with the CpHMD simulations as benchmarking, and discuss the somewhat unusual titration of several acidic residues. The most widely used pKa prediction methods, though useful for globular proteins, fail to capture the specificities of channel proteins embedded in biological membranes. This is the first attempt we know to use CpHMD to study the pH-dependent charge of a large multiionic channel (with over three hundred ionizable residues) embedded in a lipid membrane.

## Introduction

Electrostatic interactions play a key role in protein structure and function (1, 2). Protein folding and ligand binding as well as many biological processes like transport of ions and small metabolites across cell membranes or enzyme catalysis are tightly regulated by the solution pH. This fact evidences the key role of the protonation or deprotonation of the protein ionizable residues. X-Ray crystallography, Nuclear Magnetic Resonance (NMR) and Cryo-Electron Microscopy have allowed obtaining the 3D structure of an increasing number of proteins at atomic resolution. However, this high-resolution structural information needs to be supplemented by the pH-dependent charge state of the protein amino acids, often quite different from what would be expected in a free solution because of the protein low dielectric environment and the mutual interaction between ionizable sites. In addition, the local environment of the ionizable residues is highly dynamic and there is a mutual influence between their protonation state and their structural conformation. This is even clearer in protein channels because of the hydrophobic (low polarizability of the membrane) or hydrophilic (higher polarizability in the aqueous pore) environment surrounding the protein residues depending on their location. Also, the net charge and dipole moment of the membrane polar heads may influence the pKa of the residues in their vicinity (3). These facts have motivated the great effort made in the experimental and theoretical characterization of the ionization equilibria in those sites, that is, in measuring (4, 5) or theoretically predicting their pKa.

There are a relatively large group of protein channels whose ion selectivity, conductance and gating are largely controlled by the protein electrostatic properties while ion specific transport properties play a minor role. This is the case of general diffusion porins and some toxins, which are sometimes included in the family of unconventional channels (6). Even in non-specific channels the charge state of the protein amino acids is a key factor not only in the channel discrimination of small inorganic ions (7), but also in antibiotic translocation (8) and in their use as analyte biosensing nanodevices (9, 10).

Here we study the charge state of the outer membrane porin F (OmpF) from Escherichia coli. This channel is a major pathway for small hydrophilic molecules through the outer cell wall. The crystal structure of OmpF was resolved more than thirty years ago (11), and there is a large body of experimental characterization (conductance, selectivity, gating, fluctuation analysis, antibiotic permeation, etc.) on the wild-type and mutants (12–24) of this beta-barrel protein. The importance of this general diffusion porin lies in the fact that it has been identified as one of the key pore-forming proteins involved in antibiotic translocation across the outer membrane of E. coli and other Gram-negative bacteria (8, 25, 26). Matching between the charge state of key channel residues and the charge of permeating antibiotic molecules is crucial for antibiotic resistance (27, 28). In addition, modeling and simulation of ion and small solutes transport across this channel requires prior knowledge of the ionization state of amino acid residues and the protein dielectric constant (as well as its value in the aqueous pore and in the membrane). However, the set of local charges and the dielectric constant are not independent from each other. In fact, the dielectric environment regulates the ionization of neighbor residues (29, 30). In absence of experimental measurements of ionization constants of OmpF titratable groups (as happens with most protein channels), several pKa prediction methods based on an implicit representation of the protein, the membrane and the solvent and the use of the linearized Poisson-Boltzmann (PB) equation (15, 31–33) have been used. Interestingly, all MD simulations of OmpF channel performed over almost twenty years have used nearly the same charge state at neutral pH, despite notable advances in the pKa prediction methods (32–37).

All Continuum electrostatics methods employed for pKa calculation face the same problem: they need to assume a given value for the dielectric constant of the protein, which is unknown a priori. Many authors have discussed the best assumption of dielectric constant to yield good agreement with experimental results. As pointed out by Varma and Jakobsson (33), this question remains unanswered possibly because of the impossibility of describing the complex dielectric properties of the membrane by a single dielectric constant. Furthermore, the linear addition of electric potentials arising from neighbor charges (an assumption not valid when using the full nonlinear PB solution) is also a challenge of continuum electrostatics methods. Recently, other high-throughput methods for protein pKa prediction have become popular which can be broadly catalogued as empirical and MD-based.

Empirical methods, such as PROPKA (38, 39) and DeepKa (40, 41), estimate pKa values based on statistical models that correlate structural features of proteins with known pKa values. These methods are popular due to their speed and simplicity, providing reasonably accurate pKa estimates for many residues, mainly in relatively stable environments. However, empirical methods tend to overlook the dynamic nature of proteins, particularly in complex environments like membrane channels, where conformational changes and interactions with surrounding water molecules significantly impact protonation states. As a result, empirical methods may overestimate the protonation of residues embedded deep in the protein or within highly dynamic regions, where solvent accessibility and local conformational shifts are crucial.

PROPKA calculates the pKa of ionizable residues in a protein using an empirical and physical rule-based approach. It fine-tunes its calculations using a dataset of experimental pKa values for various residues in different protein environments. DeepKa uses deep learning to process structural and environmental features to predict the pKa. While PROPKA uses a dataset of measured pKa, DeepKa is trained on pKa computations. Both methods have the advantage of a fast, often reliable, pKa prediction.

Constant pH Molecular Dynamics (CpHMD) offers a more advanced approach by simulating the protonation states of residues in a dynamic, time-dependent manner (42). Unlike empirical and PB-based methods, CpHMD allows the protonation states of residues to fluctuate in response to real-time environmental interactions, such as local conformational shifts, water dynamics, and electrostatic changes. This approach is particularly valuable for studying membrane proteins like the OmpF channel, where residues experience varying degrees of exposure to the solvent and interact with other charged residues in a dynamic environment. In Gromacs CpHMD, this method is implemented in a way that integrates the protonation dynamics with the conformational flexibility of the system, enabling the simultaneous observation of structural and protonation changes. It uses a molecular dynamics scheme based on λ-dynamics in which a λ-coordinate is introduced for every titratable residue and integrated together with positions of regular atoms (43, 44).

Here we use CpHMD to calculate the pKa of acidic residues of OmpF channel (aspartates and glutamates) that might be a priori responsible for the changes in the channel charge state upon titration from pH 8 down to pH 1. These pKa are compared to those previously reported by a PB-based approach (15), a recent popular PB-based pKa predictor, H++ (45) and with values obtained from the two abovementioned empirical methods. We discuss the differences in the pKa yielded by these five methods and the large pKa shifts predicted for certain residues as well as the implications of their titration curves. Some large pKa shifts suggest strong interactions between neighbor residues. We also discuss the apparent negative cooperativity that stems from some titration curves that do not follow the Henderson-Hasselbalch equation. This analysis of the entire titration curve cannot be done with any of the empiric methods, which only yield pKa predictions. We show that such apparent negative cooperativity exhibited in a few cases may be the result of microstate heterogeneity of some residues that, at least in one monomer, explore more than one conformation.

## Methods

### System setup

The PDB structure of the OmpF protein 2OMF (11) was used as starting model. The initial system was prepared using the CHARMM-GUI membrane builder (46). The OmpF protein was embedded in a bilayer of 595 molecules of 1,2-Dipalmitoyl-sn-glycero-3-phosphocholine (DPPC). Both embedding and subsequent solvation in a box of 15.0 × 15.0 × 15.8 nm^3^ were performed using the CHARMM-GUI membrane builder. The CHARMM36-mar2019-cphmd force field (47) and CHARMM TIP3P water model were used for topology generation. Here, cphmd signifies the inclusion of CpHMD-specific modifications of bonded parameters for the titratable versions of Asp, Glu, His, Arg and Lys. Details on these modifications are described in (44). The simulation input files were set up using an in-house python script: initially, all Asp, Glu, His, Arg, and Lys residues were made titratable. CpHMD parameters for Asp, Glu, Lys, and His were obtained from (44), while parameters for Arg were obtained from (48). We run several simulations at different pH ranging from 1 to 10 and found that only acidic residues (Asp and Glu) change their protonation state, while the basic residues remain in their default protonated state during all the procedure. All the remaining simulations were then performed with only acidic residues allowed to change protonation state, which resulted in an increased computational performance. After selection of the titratable residues, KCl was added to ensure a net-neutral system at *t* = 0, and to establish an ion concentration of 150 mM. Additionally, 200 buffer particles were added to compensate for charge fluctuations and maintain a net-neutral system at *t* > 0. For more information on the buffer particles, refer to (43, 49). Finally, an in-house python script was used to generate all CpHMD-specific GROMACS input files.

### CpHMD simulations

CpHMD is based on the λ-dynamics technique developed by Brooks and co-workers (50). A one-dimensional λ-coordinate with fictitious mass m_λ_ was introduced for each titratable site, and the equations of motion for these additional degrees of freedom were integrated along with the Cartesian positions of the atoms (51). All CpHMD simulations were run using the GROMACS CpHMD beta. This version is based on the 2021 release branch and modified to include the routines required for performing the λ-dynamics calculations. The source code branch is maintained at www.gitlab.com/gromacs-constantph until it has been fully integrated into the main distribution. Energy minimization used the steepest-descent algorithm. Relaxation was initially performed in the NVT ensemble for 250 ps with a time step of 1 fs, using the Berendsen thermostat (52) with a coupling time of 1 ps and a temperature of 300 K. Bond lengths were constrained using the LINCS algorithm (53), and electrostatics were performed using the PME method (54). Subsequent relaxation and production runs were made in the NPT ensemble using a time step of 2 fs, with pressure kept at 1 bar using the Berendsen barostat (52) (coupling time 5 ps) while gradually releasing restraints on the heavy atoms. Subsequently, three independent runs of 200 ns were performed with the restraints fully released and the CpHMD λ-dynamics activated. We modified the barrier of the residue D312 (in the three monomers) from the default value of 5 kJ·mol^-1^ to a value of 16.5 kJ·mol^-1^ as described elsewhere (43). Assuming the three OmpF monomers are structurally identical, the averaging of the protonation states is made over a total equivalent time of 1.8 µs.

### Analysis

Mean protonation fractions were obtained by averaging the time-averaged protonation fraction values from the three monomers across the six replicates for each system. Protonation fractions were defined as Nproto/(Nproto + Ndeproto), where Nproto and Ndeproto are the number of simulation frames in which the residue was considered protonated (λ < 0.2) or deprotonated (λ > 0.8), respectively. We also determined protonation fractions using a simple averaging over the lambda variable. The differences in all cases between both methods were well below 1%, so that we used the latter for the results presented here. Python scripts (55) were used extensively for trajectory analysis, which included representation of titration curves, determination of the pKa of each residue, and generation of summary files in spreadsheet files. The pKa’s were determined by finding the pH at which the protonation fraction was 0.5, by using a linear interpolation between the range of discrete pH values explored in CpHMD simulations. In this work we used pH values ranging from 1 to 8 in increments of a pH unit. For residues with titration curves following sigmoid-like HH equation, we additionally fitted the protonation fraction values to the HH model, using the pKa as a fitting parameter. Differences between the pKa values from both procedures were not significant (On average less than 0.04 pKa units). We used GromacsWrapper (56) for the reading and processing of the resulting xvg files from the gmx cphmd command. Protein renderings were generated using Pymol (57).

## Results and Discussion

### Comparison between pKa predictions using different methods

We analyzed the acidic residues of OmpF channel (aspartates and glutamates) that might be relevant to the changes in the channel charge state upon titration from pH 8 down to pH 1. The choice of this pH range obeys to the fact that E. coli can withstand very acidic environments *in vivo*, like that in the extremely acidic stomach (pH range 1.5– 3.5) (58, 59). We compare the OmpF pKa values reported by Alcaraz et al. (15), who used a PB solver of UHBD (hereafter denoted as PB_A), the predictions provided by the pKa predictor H++ (45) (also based on classical continuum electrostatics), the values provided by the empirical pKa predictors PROPKA (38, 39) and DeepKa (60), and those yielded by our CpHMD simulations in 150 mM KCl solutions when the channel is embedded in a neutral DPPC membrane. Additional CpHMD simulations involving the basic residues (arginines and lysines) were also performed (not shown here), which showed that the charge state of these basic amino acids remains unaltered within this pH range of 1 to 8.

Figure 1 displays the pKa calculated according to the five methods above mentioned together with the model pKa (61) depicted as a reference dash line. Glutamates (panel A) and aspartates (panels B and C) are shown separately. There are another 7 residues that are always protonated (pKa > 8) or deprotonated (pKa < 1) in this pH range according to the CpHMD prediction. These are later analyzed separately and are not included in Figure 1. The actual pKa values of all the residues predicted by the five methods are listed in Table S1.

**Figure 1.**
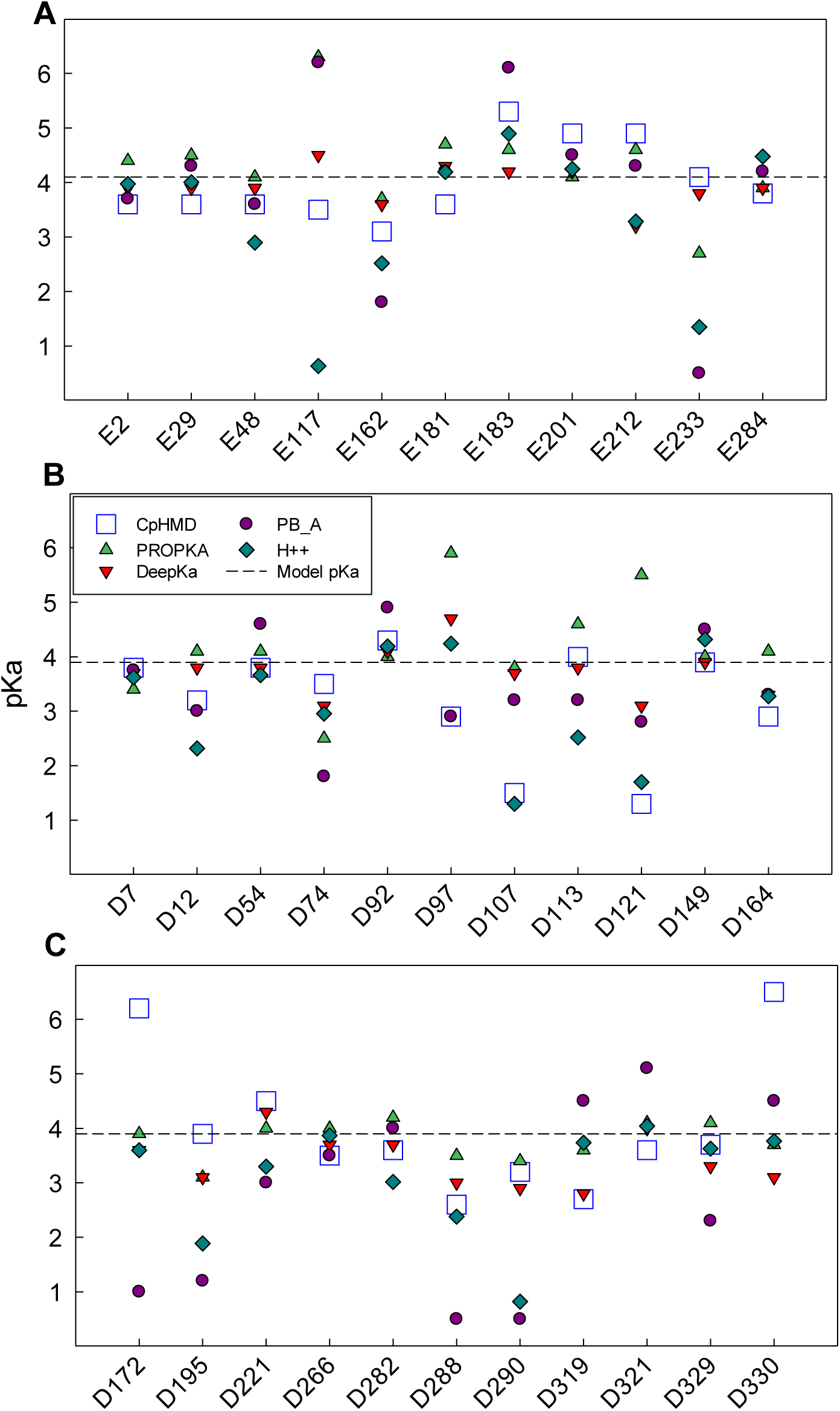
pKa prediction of the five analyzed methods for each acidic residue in OmpF. Glutamates are shown in panel **A,** and aspartates in **B** and **C.** Residues with anomalous ionization (pKa < 1 or pKa > 8) are not shown here. Model pKa is depicted by a dash line.

The first feature that stands out is that there are positive and negative pKa shifts among the predictions of all methods. If the main factor in the deviation of the pKa of a residue from its model pKa were the low polarizability of its environment, the pKa shifts should be positive since, in general, the residues buried in the protein have a higher probability of being protonated. As can be seen in Figure 1, this trend is not general, which suggests that in many cases the energy of electrostatic interaction with neighboring residues is high enough to compensate for the Born or solvation energy penalty. Actually, two thirds of the pKa shifts predicted by our CpHMD simulations are negative.

Second, we observe that for a small number of residues, the pKa predictions of the different methods diverge significantly. This is the case for a key residue in the OmpF constriction: E117 (with predicted pKa values between 0.6 and 6.3) and other less significant residues in the channel such as E233, D121 and D172.

The two methods based on PB electrostatics differ considerably in their pKa shift prediction (Figure 2A). The RMSD between them is ca. 1.4 pKa units. This might seem surprising, given the similar approach of both methods. However, the evolution from UHBD (1991) to H++ (2005) demonstrates notable advancements in computational methods for pKa prediction. UHBD, which solves the PB equation using a finite-difference approach, provides detailed electrostatic potential calculations and global charge distribution analysis but assumes a rigid protein structure, limiting its ability to capture dynamic effects such as conformational changes linked to protonation. In contrast, H++ offers a more flexible approach by incorporating two methods for pKa prediction: a clustering algorithm for systems with fewer than 80 ionizable residues, as in the case of OmpF (41 acidic residues per monomer), and Monte Carlo simulations (like traditional PB methods) for larger systems. H++ also introduced side-chain conformational sampling for histidine, glutamine, and asparagine residues, which, while not directly relevant to acidic residues like aspartates and glutamates, could influence their protonation states through neighboring interactions. This ability to model localized conformational flexibility makes H++ better suited than UHBD for capturing protonation-coupled dynamics in protein channels, such as the behavior of acidic residues in the constriction zone of OmpF. Overall, H++’s flexibility, speed, and structural adaptability provide a more accurate and practical solution for pKa predictions in dynamic systems like protein channels, where localized and neighboring effects play critical roles.

**Figure 2.**
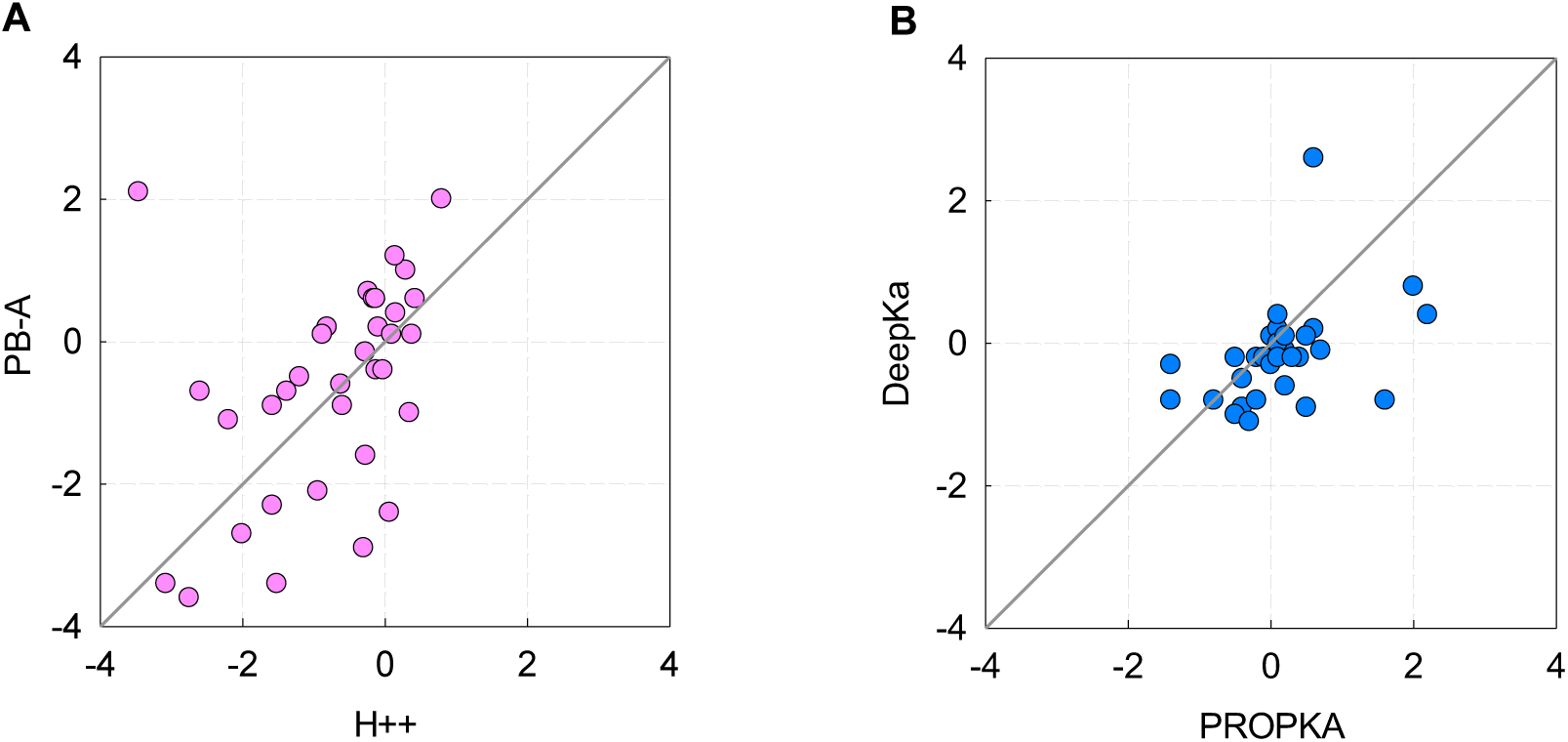
pKa shift prediction of the two methods based on PB electrostatics (panel **A**) and the two heuristic methods (panel **B**). pKa shift mean deviation with respect to the model pKa.

The comparison of pKa shifts predicted by the two heuristic methods, PROPKA and DeepKa (Figure 2B) also reveals some differences between the two methods, although not so large as between the two PB-based methods. The RMSD between them is 0.8 pKa units. The key to these differences must be sought in the set of experimental measurements (PROPKA) or protein pKa calculations (DeepKa) used in the training of each predictor. In addition, generally both methods predict lower pKa shifts than the two PB based predictors.

Table 1 summarizes the performance of PROPKA, DeepKa, PB_A, and H++ in the pKa prediction of acidic residues (Asp and Glu) by taking CpHMD as benchmarking. It displays the overall Root Mean Square Deviation (RMSD) of pKa from each method with respect to CpHMD calculation, which is reportedly the most reliable pKa prediction (40).

**Table 1.** Overall pKa RMSD.

| **pKa predictor** | **RMSD from CpHMD** |
| --- | --- |
| DeepKa | 0.96 |
| PROPKA | 1.20 |
| H++ | 1.38 |
| PB_A | 1.83 |

The comparison between the two empirical predictors, DeepKa and PROPKA, shows that DeepKa outperforms PROPKA in terms of lower RMSD values. DeepKa achieves the lowest RMSD among all methods, aligning with the findings of Wei et al. (62), where DeepKa presented the best results among several pKa predictors analyzed. PB_A exhibits the highest RMSD (∼1.83), indicating significant deviations from reference (CpHMD) pKa values. Its performance is consistent with the challenges of PB-based approaches. H++ demonstrates intermediate performance, with deviation metrics falling between those of PROPKA and PB_A. While not as accurate as DeepKa, it shows a relatively good agreement with CpHMD data.

The observation that DeepKa predictions yield lower RMSD values is not surprising, given that DeepKa was trained using data derived from CpHMD calculations, and thus a better agreement between the two methods could be anticipated. The consideration of CpHMD as the method of better performance, which justifies its selection as a reference in DeepKa, is based on theoretical arguments, as this methodology more accurately represents both the structure and dynamics of the protein along with the various interactions present in the environment of the residues. Additionally, this superiority is also supported by experimental data, where CpHMD has outperformed other methodologies in reproducing experimentally measured pKa values (40).

On the other hand, although the overall agreement between CpHMD and DeepKa predicted pKa values is not bad (Table 1), significant divergences are observed for certain residues, particularly D256 and to a lesser extent E71, D127 and D330. An analysis of the position of these residues within the protein environment reveals that some of them are located close to the protein-lipid interface. Given that DeepKa was trained and validated using water soluble proteins (40, 41), it is expected to yield a poorer prediction for residues that do not meet this condition.

### Residues with anomalous ionization (deprotonated or protonated over the entire pH range 1-8)

Some acidic residues exhibit anomalous ionization according to CpHMD simulations, remaining deprotonated (pKa < 1) or protonated (pKa > 8) across the pH range studied. D37, E62, E71 and D126 stay charged for the entire pH range, whereas D127, D256, E296 and D312 stay in their neutral form. This prediction contrasts with that of the other four methods explored. The pKa obtained using empirical methods show the greatest differences, while methods based on the PB equation agree in some cases with CpHMD (see Table 2). To find an explanation, we examined the local environment of each residue, including nearby titratable residues, interactions with water molecules, and proximity to the monomer-monomer interface (Figure 3).

**Figure 3.**
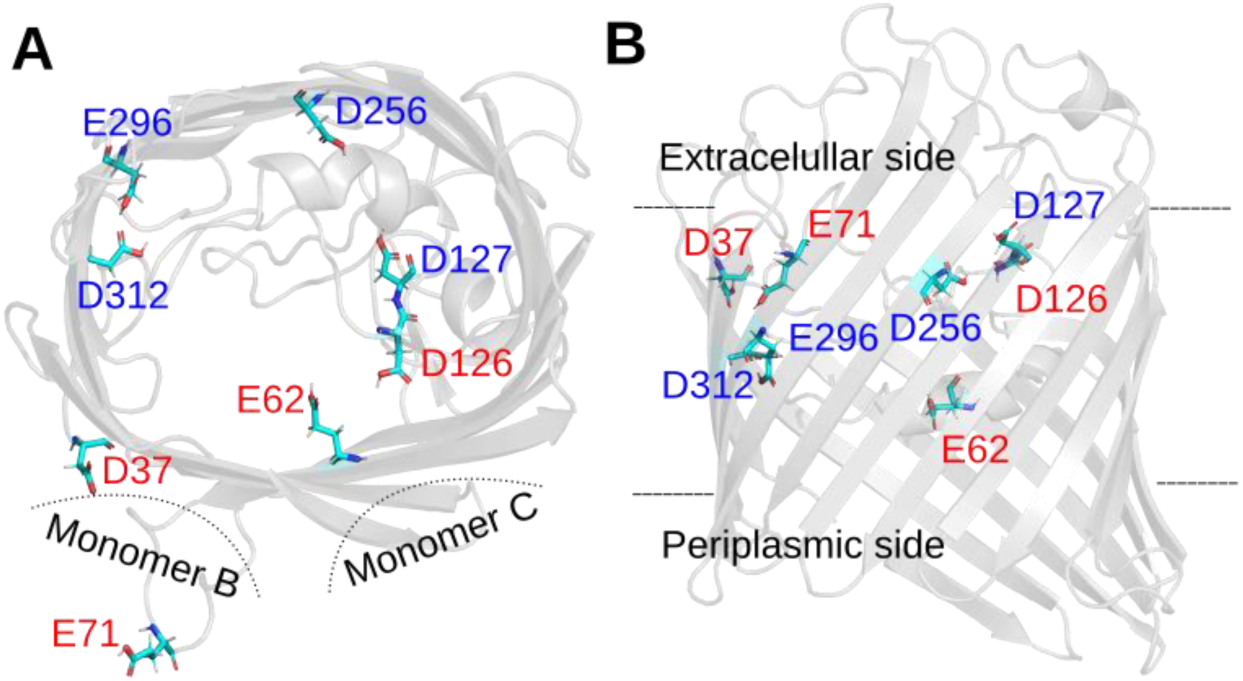
Representation of OmpF acidic residues with anomalous pKa values. The four residues that remain deprotonated across pH 1-8 are red labeled and the four residues that are protonated in the same pH range are blue labeled. A) Top view of OmpF chain A. B) side view (90° rotation around the x-axis). Dash lines show the relative position of the other monomers (in panel A) and the approximate limits of the membrane (in panel B).

**Table 2.** pKa values of residues with very large pKa shifts.

| Residue | CpHMD | PROPKA | DeepKa | PB_A | H++ |
| --- | --- | --- | --- | --- | --- |
| D 37 | < 1 | 4.9 | 3 | 0.3 | 2.9 |
| E 62 | < 1 | 2.5 | 2.6 | 0.4 | < 0 |
| E 71 | < 1 | 1.1 | 4.1 | 0.5 | 4.6 |
| D 126 | < 1 | 3.2 | 2.5 | 0.5 | < 0 |
| D 127 | > 8 | 7.4 | 5.5 | 3.8 | 6.6 |
| D 256 | > 8 | 6 | 3.2 | 0.6 | < 0 |
| E 296 | > 8 | 9.6 | 7.3 | 8.9 | 9.4 |
| D312 | > 8 | 4.5 | 6.5 | 1.5 | 4.0 |

The local environment of D37 likely favors its deprotonated (charged) state. Although D37 is located at the monomer-monomer interface of the OmpF trimer, this region is still water accessible. Moreover, the interaction with nearby residues such as His21 and Tyr22 may favor the deprotonated form through hydrogen bonding or dipolar interactions, which can help accommodate the negative charge by reducing the energetic cost of maintaining a charged carboxylate in this region. If His21 is protonated, its positive charge may contribute to a local electrostatic potential that favors the anionic form of D37 and creates a barrier to proton access, further lowering its pKa. Even in a neutral state, His21 may still participate in dipolar interactions or hydrogen bonding with D37, which can also contribute to its deprotonation.

The protonation state of E62 is shaped by a network of nearby residues located at the OmpF monomer-monomer interface, including Tyr14, Lys16, Arg42, Lys46, Arg82, and Tyr102. The cluster of basic residues, lysines and arginines, creates a positively charged environment that favors the deprotonated form of E62. As a result, its pKa can shift significantly downward compared to its intrinsic value in solution. Although this region is involved in monomer-monomer contacts, it remains accessible to water. The large negative pKa shift reflects the dominant influence of these surrounding basic residues.

E71 experiences a weak direct influence from neighboring residues belonging to the same monomer –limited primarily to D74– but due to its position at the monomer-monomer interface (see Figure 3), this is compensated by a cluster of basic residues from the adjacent monomer, including K80, R82, and R132. Their collective electrostatic influence may favor the deprotonated form of E71, keeping it negatively charged down to pH 1 and below.

D126 is surrounded by a dense cluster of positively charged arginines –R100, R132, R163, R167, and R168– which collectively exert a strong effect favoring its deprotonated form. The nearby titratable residue D127 provides a counteracting effect, but not big enough to compensate for the positive charges. D126 and these arginines are located within the monomer–monomer interface, a relatively restricted region where their side chains are oriented internally facing each other. This close arrangement of positive charges likely creates a strong local electrostatic field that hamper proton association. At the same time, the limited space in this interface may hinder solvent access, making it physically more difficult for a proton to reach D126. As a result, D126 remains deprotonated across the studied pH range.

The protonation state of the aspartate D127 is modulated by nearby acidic residues D126 and D256 as well as basic arginines R132, R167, R168, and R196. Structurally, D127 is in a partially buried region, with low solvent exposure. Interestingly, earlier structural and biochemical studies have highlighted D127 unusual behavior. Karshikoff (31) observed in the OmpF crystal structure that the carboxylate side chain of D127 is positioned within hydrogen-bonding distance (0.26 nm) of the backbone carbonyl of residue A237, a spatial configuration that supports a protonated state. However, subsequent MD simulations by Varma et al. (33) suggested that both protonated and deprotonated forms of D127 are energetically feasible, depending on the local dielectric environment. Experimental work using cysteine-substitution mutants also revealed that D127 is poorly accessible to solvent, implying that the residue is –at least partially– buried (63). While these earlier findings are not conclusive about D127 charge state under physiological conditions, our CpHMD data support a persistently protonated form, consistent with a buried, hydrogen-bond-stabilized environment.

D256 experiences multiple favorable interactions that support protonation, including contributions from neighboring residues such as D121, Y124, D127, Y231, and E233. The local negative electrostatic potential likely lowers the energetic cost of protonation, thereby favoring the protonated state. Structurally, D256 is in a region near the lipid interface –far from the adjacent monomers– with its side chain oriented internally; all key interacting partners are also in the same monomer.

E296 position in the monomer is like that of D256. It also remains protonated between pH 1 and 8 in our CpHMD simulations, in agreement with earlier computational studies (31, 33). E296 protonation state is influenced by moderate contributions from nearby residues, including E117, D121, Y294, and Y310, and shows particularly strong coupling with D312. All these residues belong to the same monomer. Earlier work by Varma and Jakobsson (32) highlighted the presence of a protonation-coupled network involving E296, D312, Y22, Y310, and E117, where shifts in the protonation state of one residue propagate through the network, altering the charge distribution and potentially the conformational dynamics of the protein. Pongprayoon (64) suggested that full deprotonation of both E296 and D312 significantly increases OmpF flexibility, particularly in the loop L3 region, highlighting the functional sensitivity of the channel to the charge states of these residues. Our CpHMD simulations predict that both E296 and D312 remain neutral from pH 10 down. Under these conditions, we observe no substantial increase in loop L3 flexibility, nor any significant changes in the distances between E296/D312 and E117—the closest residue in loop L3. Pongprayoon (64) also reported that when both residues are charged, the constriction radius decreases. This behavior is not observed in our results when comparing the constriction radius from the CpHMD- derived structure with that from the crystallographic PDB structure. It is worth noting that in Pongprayoon’s study the protonation states of E296 and D312 were manually assigned, without accounting for possible effects on neighboring residues.

### Other residues with large pKa shifts: D107 and D121

D107 and D121 exhibit considerably larger pKa shifts than most acidic residues. Their predicted pKa are 1.5 and 1.3, respectively. Both residues are near the OmpF constriction zone (Figure 4). In fact, D121 is reported by several authors as one of the acidic residues of the constriction, positioned on the loop L3 which is key in the regulation of the channel ion conduction. The simplest explanation of their big negative pKa shift is their positively charged environment. D107 is near arginine R140, while D121 is near the group of positive charges, K80, R132, R168 and R167. These positive charges might increase the energetic barrier for protons to bind both aspartates, thus decreasing their pKa’s.

**Figure 4.**
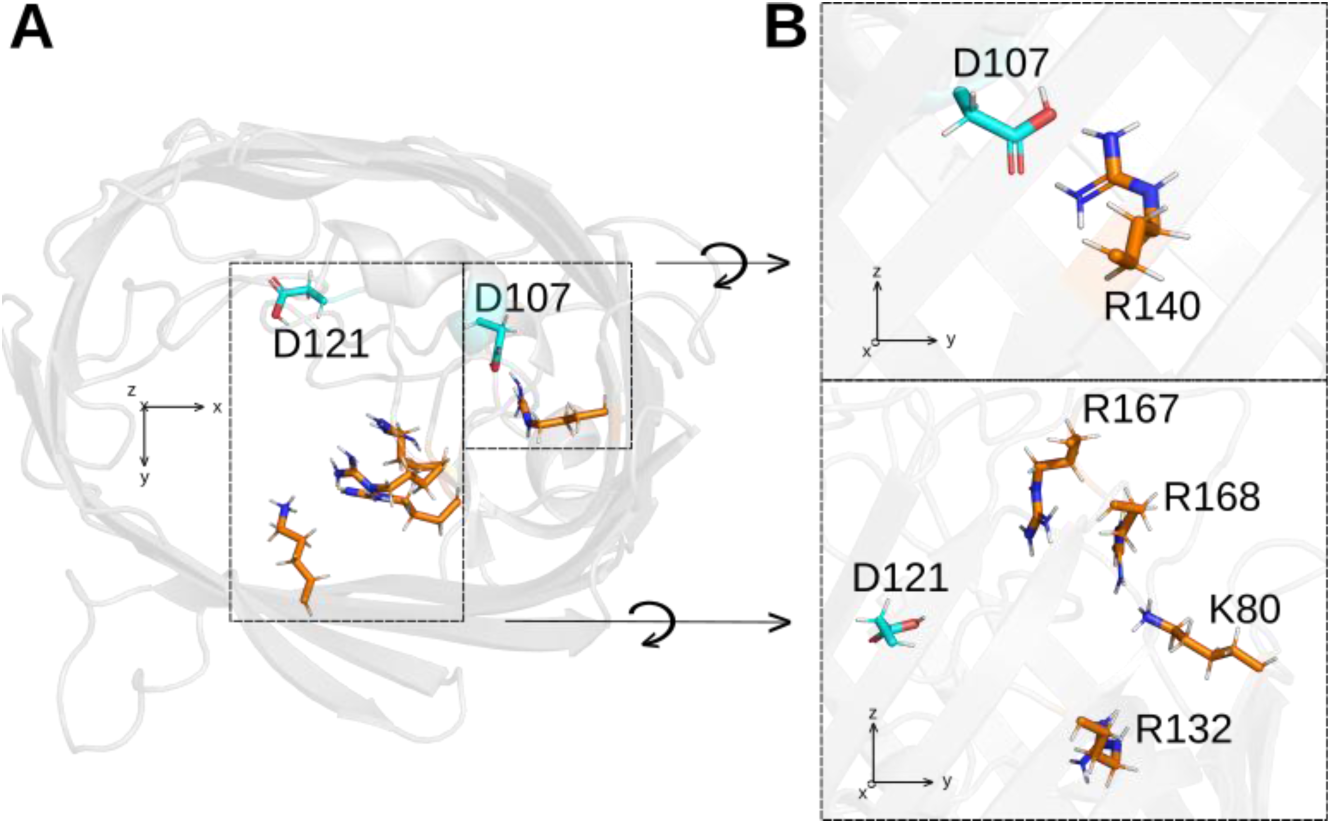
Representation of D107 and D121 and the cluster of basic residues supposedly responsible for their large pKa shift. Top view in panel **A.**, and side view in panel **B.**

It is worth noting that unlike the transport of small inorganic ions, in which channel constriction plays a predominant role, antibiotic permeation involves other ionizable amino acids from the periplasmic and extracellular vestibules. The study by Ziervogel and Roux (65) shows that two of the residues that we have identified with anomalous ionization (D121 y E62) stabilize the binding of ampicillin and carbenicillin positively charged NH3+ moieties, respectively, to OmpF channel. More recently, Acharya et al. (27) also reported that the permeation of charged antibiotics may involve interactions with the L3 loop, particularly with residue D121, further supporting its critical role in antibiotic translocation.

### Residues displaying “slow” titration

Given the number of titratable sites in the OmpF protein close to each other, one would expect a priori that their ionization should be far more complex than that of a collection of independent sites with titration curves following the standard HH sigmoidal shape. For small deviations from HH equation, titration curves are commonly fitted to Hill equation (66), Eq. (1), where *n* is the Hill coefficient, a measure of the cooperativity in a ligand binding process (the protonation of an acidic residue in our case). Then, the fraction *θ* of identical sites that are protonated (or the protonation probability of a single site) is given by

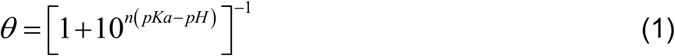

If a residue charge state is not affected by the protonation of its neighbors, its titration curve follows the standard HH equation, i.e., *n* = 1. But, in the opposite case we have two possible scenarios: *n* > 1 or *n* < 1, which represent positive and negative cooperativity, respectively.

We averaged the charge state of each one of the 41 acidic residues obtained in the three replicas of the three OmpF monomers and fitted them to a) the HH equation to obtain the pKa and b) to Hill equation (taking pKa and *n* as free parameters). In most cases the two nonlinear fittings yielded virtually the same pKa (differences less than 0.1) and values for *n* very close to 1. However, there were a few exceptions in which the average of the 9 protonation curves (3 replicas x 3 monomers) was much better fitted to Hill equation (with *n* values around 0.3-0.4) than to HH equation. Also in these few cases, both HH equation and Hill equation led to virtually the same pKa. A priori, values of the Hill coefficient lower than 1 are associated to negative cooperativity between neighbor titratable sites, i.e. shallower titration curves, which implies strong interaction between them. However, negative cooperativity could be also the result of heterogeneity of microstates of a given residue (67, 68). There exists the possibility that a residue spends some time in a spatial conformation (characterized by a local minimum of free energy) with a corresponding pKa that differs from the most probable conformation pKa. If that were the case, the process of averaging of all protonation states along the MD trajectory might yield an apparent negative cooperativity without true physical interaction meaning. Figure 5 shows as an example the protonation state of D97 along an extended pH range. As seen, fitting to Hill equation is better than to HH equation. The pKa obtained by interpolation (the pH at which *θ* = 0.5) is 3.0, not very different from the best fitting values according to HH or Hill equation (3.1 and 3.3, respectively).

**Figure 5.**
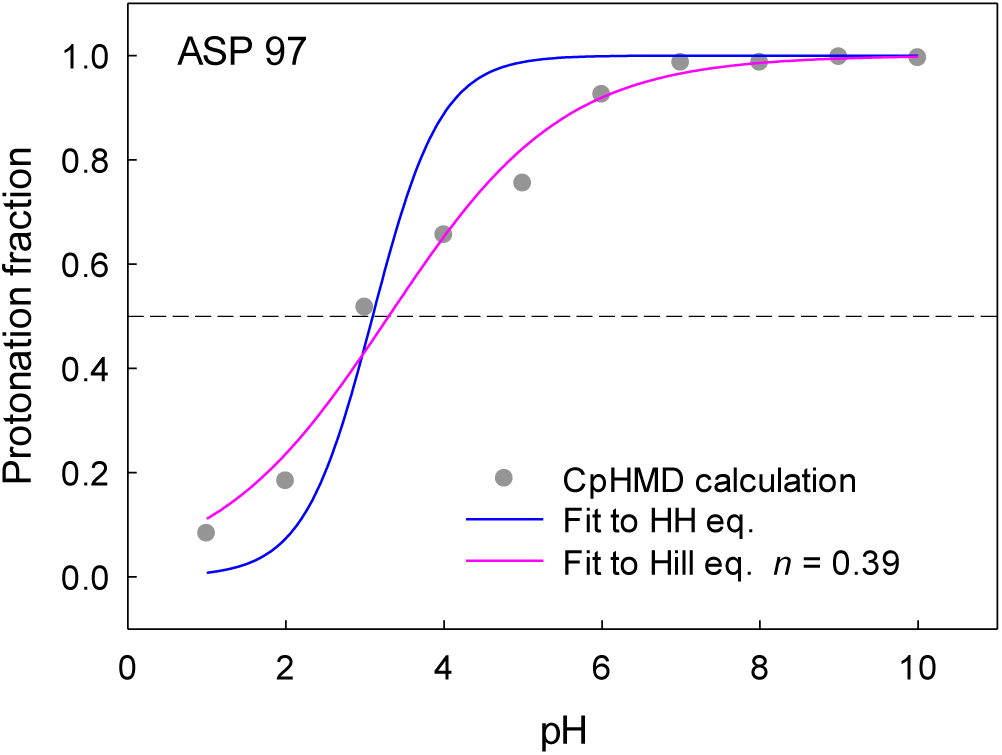
D97 is an example of OmpF acidic residues that display shallower titration curves. The average fraction of protonation is displayed together with the best fitting curve according to HH equation (blue) and according to Hill equation (pink) and the fitting parameters (pKa and Hill coefficient *n*).

By looking at the protonation states and RMSD of D97 at pH 4 (Figure 6), we see that this residue is accessing different microstates along the simulation time. For instance, during a relatively long interval between 100 ns and 175 ns, monomers #1 and #3 display almost no difference between them in the three replicas (not shown) but in monomer #2 D97 exhibits different protonation states in the three replicas (Figure 6A). In the third replica D97 is protonated all the time (100-175 ns) while in the first and second replicas the residue alternates between its charged and uncharged state (what would be expected for pH 4, not far from the pKa 3.3). Interestingly this “unexpected” protonation state seen in the third replica and lasting ca. 80 ns is mirrored in the RMSD of D97 (Figure 6B) within the same time interval. In addition, this “unusual” protonation state in the third simulation replica of monomer #2 might be related with a conformation change of D97 and its closest neighbor residues, Y58 and K89. Figure 6C shows a slight change in the orientation of D97 and K89 in the third replica. Tyrosine Y58 is not titratable and remains with the same orientation in the three replicas for the time interval considered. It is plausible that this and other possible microstates of D97 contribute to the averaging of the simulation trajectory by slowing down the titration curve but with minimal or no effect on the calculated effective pKa of the residue.

**Figure 6.**
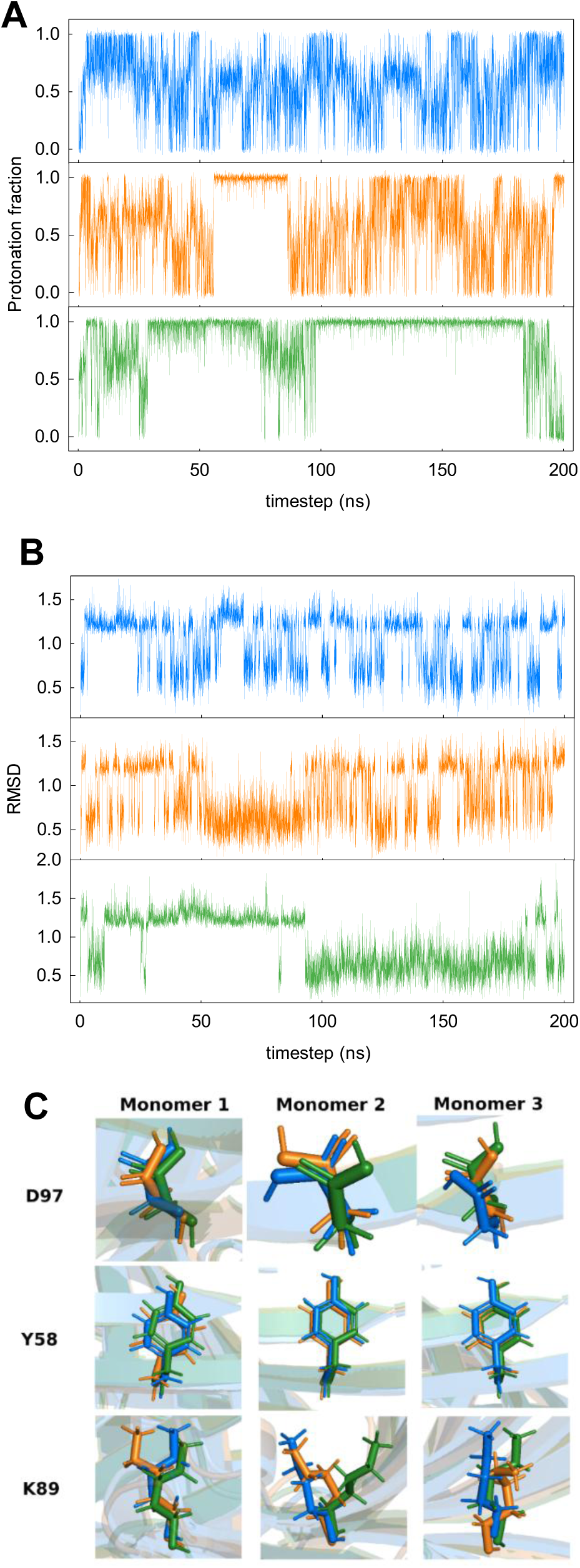
Protonation states (A) and RMSD (B) of monomer #2 along the simulation time. Representation of D97 side chain and its closest neighbors Y58 and K89 (C). Frame corresponding to 150 ns. The three replicas are colored blue, orange and green, respectively.

### OmpF net charge in acidic environments

The differences between the pKa predictions obtained from each method become somewhat blurred when calculating the overall net charge of an OmpF monomer. This is a consequence of the positive and negative pKa shifts yielded depending on the residue. However, as shown in Figure 7, some general conclusions can be extracted. Firstly, all apparent titration curves display a slower titration than that calculated using the model pKa of aspartates and glutamates. Fitting them to Hill equation one gets an effective pKa around 3.6-3.9 and Hill coefficient *n* around 0.5-0.7. Secondly, at neutral pH, differences in net charge calculated by different methods are very small. CpHMD predicts a total charge per monomer of −10*e*, PROPKA gives −12*e*, DeepKa yields −13*e*; PB_A and H++ predict −13*e* and −12*e*. All these values are slightly lower than the −15*e* resulting from the null model (using model pKa). As seen in Figure 7 empiric and PB-based pKa predictions overestimate the negative charge of the channel when the benchmarking reference is CpHMD. Thirdly, differences in (positive) net charge become more apparent in very acidic solutions. At pH 2, for instance, CpHMD predicts a positive net charge of 21*e*, PROPKA and DeepKa give 26*e*; PB_A yields 15*e* and H++ predicts 18*e*. All these values are slightly lower than the 28*e* resulting from the null model.

**Figure 7.**
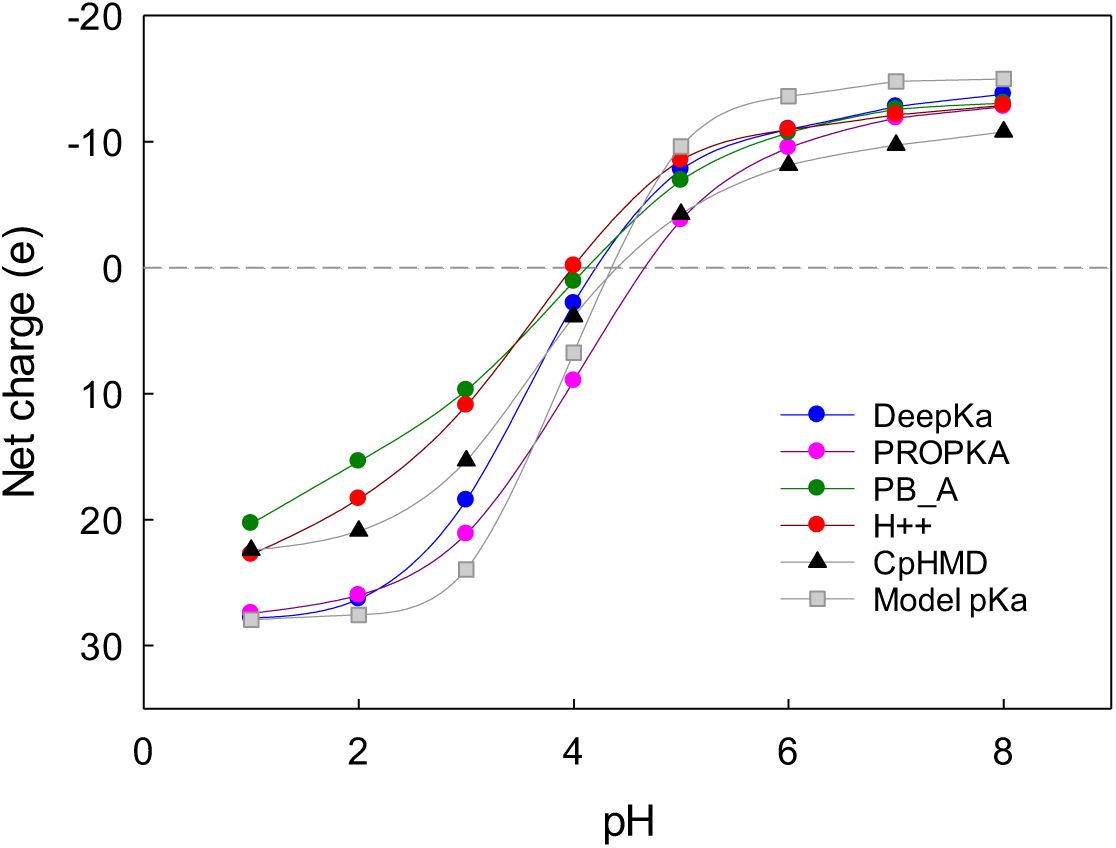
Change of OmpF net charge with pH according to each method used for calculating the protonation state of the acidic residues. The net charge per monomer includes the charge of the other ionizable residues (histidine, arginines and lysines). Solid lines are to guide the eye.

In any case, the net charge of the channel at a given pH, obtained by simply adding up the charges of the ionizable sites, does not have a direct correlation with conductance, reversal potential, or other experimentally accessible observable. Other effects must also be taken into account in the OmpF channel. First, the solvent exposed residues in the pore constriction have a key role in promoting separate pathways for cations and anions, thus enhancing the ionic conductance. In addition, the competitive binding of cations and protons to D113 and E117 also exerts a big influence on the channel permeating properties, and this process is strongly dependent on the solution pH (18, 24). Despite this, an attempt can be made to check the consistency of the pH-dependent net charge with the OmpF channel selectivity, given that it is mainly regulated by the electrostatic interaction between channel charges and permeating ions. When the channel selectivity is measured in salts with similar diffusivity of cations and anions, as happens with KCl solutions, the Reversal Potential should reflect changes in the net charge of the channel when pH titration switches the channel selectivity from cationic to anionic. Figure 7 shows the plot of OmpF Reversal Potential measured in 0.1/1 M KCl solutions at several pH (15) superimposed with the change of the CpHMD predicted net charge across the same pH range. Both datasets share a common pattern.

Figure 8 demonstrates a common pattern in the pH dependence of the net charge of an OmpF monomer, calculated by CpHMD, and the experimental reversal potential. The plot illustrates that the total charge predicted by CpHMD aligns well with the experimental reversal potential across a wide pH range, highlighting a stronger correlation in the acidic range, particularly around pH 4.

**Figure 8.**
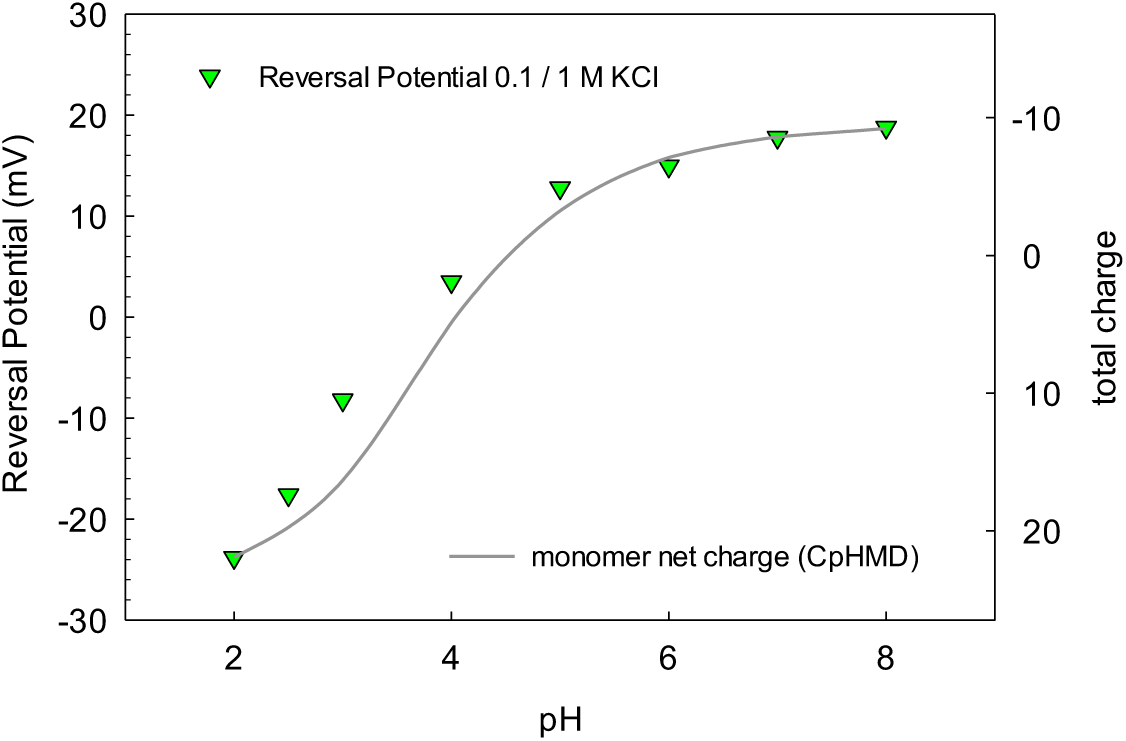
Correlation between the total charge of the monomer calculated with CpHMD (solid line) and the OmpF measured reversal potential in 0.1/1 M KCl (15) (green symbols).

This close match between CpHMD results and experimental data demonstrates the effectiveness of CpHMD in capturing the protonation states of acidic residues under varying pH conditions. The PB method tends to deviate from experimental data, especially at lower pH values. This discrepancy could be attributed to the limitations of the PB model in accounting for dynamic changes in protonation states in response to the local environment, an area where CpHMD can better capture these dynamic interactions, and these subtle shifts in charge distribution are critical for understanding the functional behavior of the OmpF channel in response to pH.

### Conclusions

Given the difficulty of experimentally measuring the pKa of channel proteins, we have resorted to calculating protonation states using one of the most advanced computational tools: the CpHMD method. Its main advantages over other pKa predictors for channel proteins can be summarized as follows. First, CpHMD captures the coupling between conformational dynamics and residue protonation. Second, it avoids the problem of a priori assigning a dielectric constant to the protein environment. This is a key challenge for pKa predictors based on PB electrostatics, which partially resolve it by assigning different polarizabilities to different protein regions depending on their exposure to the solvent or the hydrophobic core of the membrane. Finally, the CpHMD method allows the lipid membrane to be explicitly considered with its distinct polar and hydrophobic regions.

Until now, the CpHMD method has been used primarily to study pH-induced conformational changes in proton ion channels. In this work, we analyzed for the first time the pH-dependent charge of a large multiionic trimeric channel (340 amino acid residues per monomer, of which 126 are ionizable) embedded in a lipid membrane.

Comparison with other empirical methods widely used in globular proteins (PROPKA and DeepKa) and with methods based on electrostatic calculations of the static 3D structure shows significant differences in many of the acidic residues studied. In the case of the former, these differences are possibly due to the fact that the correlations and training used by both heuristic methods are based on non-transmembrane proteins. The differences with methods based on electrostatic calculations may arise from a poor representation of the local polarizability of the medium.

We have found that some residues remain unprotonated or protonated over the whole pH range [1-8]. In addition, a substantial portion of the OmpF acidic ionizable residues have a pKa different from their model pKa, and therefore their charge states must be correctly assigned. These calculated pKa may serve as a reference for future MD simulations of the channel aimed at obtaining the transport properties of both small inorganic ions and antibiotic molecules. Besides it is crucial to consider that charged metabolites crossing the channel may exert an influence on residues located near their pathway, potentially leading to changes in their pKa values and, consequently, their protonation state at a given pH. This dynamic interplay implies that pKa values alone cannot capture the full complexity of the system, and therefore, a complete CpHMD simulation would be necessary to account properly for the interaction of charged metabolites, such as antibiotics, with the channel.

The protonation states of the vast majority of 41 acidic residues of each monomer fit well the HH equation in the pH range 1–8. However, a few residues are an exception and exhibit ionization curves in which the probability of being charged varies more smoothly with pH, and the fit to the Hill equation yields a Hill coefficient around 0.3–0.4. Rather than negative cooperativity interactions between each of these residues and their neighbors, the cause of their shallower titration curves must be sought in a superposition of different microstates that differ in their pKa (67). In fact, the analysis of the MD trajectories of these residues reveals some equilibrium microstates that differ from the most probable, but of sufficient duration to influence the average of the three 200 ns replicas in each of the three monomers.

Further work awaits to study the effect of ion screening and membrane charge properties on the protonation state of the OmpF ionizable residues, as well as the role of key residues on the channel conductive and selective properties.

## Notes

### Competing Interest Statement

The authors have declared no competing interest.

### Funding

- Spanish Ministry of Science and Innovation, AEI: MCIN/AEI/10.13039/501100011033
- Universitat Jaume I: UJI-B2022-42
- Generalitat Valenciana: CIGRIS/2021/020

