## Supplemental Table S1 and Figure S1 for "Predicting residue ionization of OmpF channel using Constant pH Molecular Dynamics as benchmarking"

### Supplementary Information

#### 1. pKa prediction

Table S1 displays the pKa predictions of the 41 amino acids selected (27 aspartates and 14 glutamates) yielded by five different methods (see main text). The basic amino acids remain positively charged over the whole pH range studied [1-8].

The uncertainty associated with the CpHMD-derived values was quantified by calculating the standard deviation among the pKa values obtained from three independent replicas. This procedure involved a two-step averaging process: first, the protonation fractions were averaged over the simulated time to obtain a mean value for each residue; subsequently, these values were averaged over residues belonging to different monomers, assuming identical behavior among the three monomers. This approach effectively increases the statistical sampling, allowing for a more robust estimation of the pKa values. The resulting protonation states at each pH point yielded three titration curves, one for each replica, from which three corresponding pKa values were calculated.

Table S1

| Residue | CpHMD | PROPKA * | DeepKa § | PB_A # | H++ & |
|---|---|---|---|---|---|
| E 2 | 3.6 ± 0.0 | 4.4 | 3.9 | 3.7 | 4.0 |
| D 7 | 3.8 ± 0.1 | 3.4 | 3.7 | 3.8 | 3.6 |
| D 12 | 3.2 ± 0.0 | 4.1 | 3.8 | 3 | 2.3 |
| E 29 | 3.6 ± 0.0 | 4.5 | 3.9 | 4.3 | 4.0 |
| **D 37** | **< 1** | 4.9 | 3.0 | 0.3 | 2.9 |
| E 48 | 3.6 ± 0.0 | 4.1 | 3.9 | 3.6 | 2.9 |
| D 54 | 3.8 ± 0.1 | 4.1 | 3.8 | 4.6 | 3.7 |
| **E 62** | **< 1** | 2.5 | 2.6 | 0.4 | < 0 |
| **E 71** | **< 1** | 1.1 | 4.1 | 0.5 | 4.6 |
| D 74 | 3.5 ± 0.1 | 2.5 | 3.1 | 1.8 | 2.9 |
| D 92 | 4.3 ± 0.0 | 4.0 | 4.1 | 4.9 | 4.2 |
| D 97 | 2.9 ± 0.3 | 5.9 | 4.7 | 2.9 | 4.2 |
| D 107 | ~ 1 | 3.8 | 3.7 | 3.2 | 1.3 |
| D 113 | 4.0 ± 0.0 | 4.6 | 3.8 | 3.2 | 2.5 |
| E 117 | 3.5 ± 0.1 | 6.3 | 4.5 | 6.2 | 0.6 |
| D 121 | ~ 1 | 5.5 | 3.1 | 2.8 | 1.7 |
| **D 126** | **< 1** | 3.2 | 2.5 | 0.5 | < 0 |
| **D 127** | **> 8** | 7.4 | 5.5 | 3.8 | 6.6 |
| D 149 | 3.9 ± 0.1 | 4.0 | 3.9 | 4.5 | 4.3 |
| E 162 | 3.1 ± 0.1 | 3.7 | 3.6 | 1.8 | 2.5 |
| D 164 | 2.9 ± 0.0 | 4.1 | 3.3 | 3.3 | 3.3 |
| D 172 | 6.2 ± 0.5 | 3.9 | 3.6 | 1 | 3.6 |
| E 181 | 3.6 ± 0.1 | 4.7 | 4.3 | 4.2 | 4.2 |
| E 183 | 5.3 ± 0.1 | 4.6 | 4.2 | 6.1 | 4.9 |
| D 195 | 3.9 ± 0.2 | 3.1 | 3.1 | 1.2 | 1.9 |
| E 201 | 4.9 ± 0.1 | 4.1 | 4.2 | 4.5 | 4.2 |
| E 212 | 4.9 ± 0.4 | 4.6 | 3.2 | 4.3 | 3.3 |
| D 221 | 4.5 ± 0.1 | 4.0 | 4.3 | 3 | 3.3 |
| E 233 | 4.1 ± 0.3 | 2.7 | 3.8 | 0.5 | 1.3 |
| **D 256** | **> 8** | 6.0 | 3.2 | 0.6 | < 0 |
| D 266 | 3.5 ± 0.0 | 4.0 | 3.7 | 3.5 | 3.9 |
| D 282 | 3.6 ± 0.1 | 4.2 | 3.7 | 4 | 3.0 |
| E 284 | 3.8 ± 0.1 | 3.9 | 3.9 | 4.2 | 4.5 |
| D 288 | 2.6 ± 0.1 | 3.5 | 3.0 | 0.5 | 2.4 |
| D 290 | 3.2 ± 0.2 | 3.4 | 2.9 | 0.5 | 0.8 |
| **E 296** | **> 8** | 9.6 | 7.3 | 8.9 | 9.4 |
| **D 312** | **> 8** | 4.5 | 6.5 | 1.5 | 4.0 |
| D 319 | 2.7 ± 0.1 | 3.6 | 2.8 | 4.5 | 3.7 |
| D 321 | 3.6 ± 0.0 | 4.1 | 4 | 5.1 | 4.0 |
| D 329 | 3.7 ± 0.3 | 4.1 | 3.3 | 2.3 | 3.6 |
| D 330 | 6.5 ± 0.2 | 3.7 | 3.1 | 4.5 | 3.8 |

(*) Olsson, M.H.M., C.R. Søndergaard, M. Rostkowski, and J.H. Jensen. 2011. PROPKA3: Consistent treatment of internal and surface residues in empirical pKa predictions. *J Chem Theory Comput*. 7:525–537.

(§) Cai, Z., H. Peng, S. Sun, J. He, F. Luo, and Y. Huang. 2024. DeepKa Web Server: High-Throughput Protein pKa Prediction. *J Chem. Inf. Model*. 64:2933–2940.

(#) Alcaraz, A., E.M. Nestorovich, M. Aguilella-Arzo, V.M. Aguilella, and S.M. Bezrukov. 2004. Salting Out the Ionic Selectivity of a Wide Channel: The Asymmetry of OmpF. *Biophys. J*. 87:943–957.

(&) Gordon, J.C., J.B. Myers, T. Folta, V. Shoja, L.S. Heath, and A. Onufriev. 2005. H++: a server for estimating pKas and adding missing hydrogens to macromolecules. *Nucleic Acids Res*. 33:W368–W371.

**2. Residues whose titration curve does not fit well with the Henderson-Hasselbalch equation.**

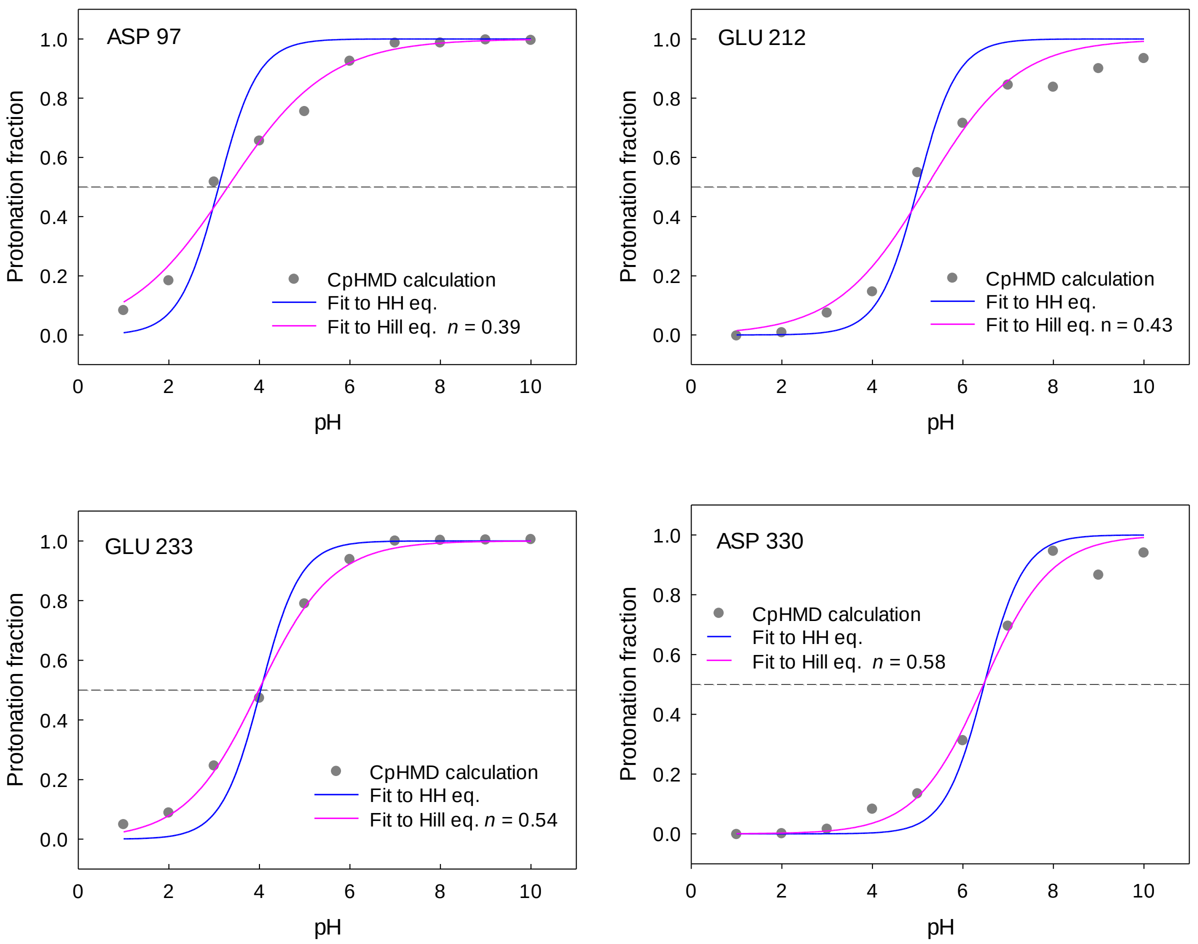


Figure S1. Titration curves of some acidic residues that depart from the standard HH equation.
